# Yersiniabactin is a quorum sensing autoinducer and siderophore in uropathogenic *Escherichia coli*

**DOI:** 10.1101/2023.02.09.527953

**Authors:** James R. Heffernan, George L. Katumba, William H. McCoy, Jeffrey P. Henderson

**Author notes:** Address correspondence to Jeffrey P. Henderson.

## Abstract

Siderophores are secreted ferric ion chelators used to obtain iron in nutrient-limited environmental niches, including human hosts. While all *E. coli* encode the enterobactin (Ent) siderophore system, isolates from patients with urinary tract infections additionally encode the genetically distinct yersiniabactin (Ybt) siderophore system. To determine whether the Ent and Ybt systems are functionally redundant for iron uptake, we compared growth of different isogenic siderophore biosynthesis mutants in the presence of transferrin, a human iron-binding protein. We observed that the Ybt system does not compensate for loss of the Ent system during siderophore-dependent, low density growth. Using transcriptional and product analysis, we found that this non-redundancy is attributable to a density-dependent transcriptional stimulation cycle in which Ybt assume an additional autoinducer function. These results distinguish the Ybt system as a combined quorum-sensing and siderophore system. These functions may reflect Ybt as a public good within bacterial communities or as an adaptation to confined, subcellular compartments in infected hosts. The efficiency of this arrangement may contribute to the extraintestinal pathogenic potential of *E. coli* and related *Enterobacterales*.

**IMPORTANCE:** Urinary tract infections (UTIs) are one of the most common human bacterial infections encountered by physicians. Adaptations that increase the pathogenic potential of commensal microbes such as *E*.*coli* are of great interest. One potential adaptation observed in clinical isolates is accumulation of multiple siderophore systems, which scavenge iron for nutritional use. While iron uptake is important for bacterial growth, the increased metabolic costs of siderophore production could diminish bacterial fitness during infections. In a siderophore-dependent growth conditions, we show that the virulence-associated yersiniabactin siderophore system in uropathogenic *E. coli* is not redundant with the ubiquitous *E. coli* enterobactin system. This arises not from differences in iron scavenging activity but because yersiniabactin is preferentially expressed during bacterial crowding, leaving bacteria dependent upon enterobactin for growth at low cell density. Notably, this regulatory mode arises because yersiniabactin stimulates its own expression, acting as an autoinducer in a previously unappreciated quorum-sensing system. This unexpected result connects quorum-sensing with pathogenic potential in *E. coli* and related *Enterobacterales*.

## INTRODUCTION

Urinary tract infections (UTIs) are one of the most common bacterial infections in clinics and hospitals, with over 7 million doctor visits and 1 million emergency room visits in the United States each year^1^. *Escherichia coli* account for ~85% of these infections and are a leading cause of recurrent UTIs^2,3^. As clinical *E. coli* isolates become increasingly resistant to first-line antibiotics, there is renewed interest in better understanding features of these and related *Enterobacterales* that increase their pathogenic potential in human hosts^4^.

Among the most prominent host-pathogen interactions in UTIs is the interplay between human factors that limit nutrient iron availability and the corresponding bacterial responses to iron scarcity^5,6^. Normal physiological iron excretion in humans is low, with most iron in the host bound to heme^7,8^. In circulation and tissue, labile iron ions are sequestered by binding to transferrin, a circulating iron transport protein that binds ferric ions in two iron binding pockets with K_a_ values of 4.7*10^20^ M^−1^ and 2.4*10^19^ M^−1^ ^9^. After the arrival of infecting bacteria, local innate immune responses in tissue intensify this iron sequestration response by introducing additional iron-binding proteins. Epithelial cells and neutrophils introduce lipocalin-2/siderocalin, which sequesters iron in complexes with catechol metabolites^10–12^. Lactoferrin, a member of the transferrin family excreted by neutrophils, can bind iron 300 times more tightly than transferrin^13^ and, like lipocalin-2, becomes detectible in the urine of patients with UTIs^14^. After bacteria establish an early infection, these innate immune responses intensify the baseline iron restriction state maintained by transferrin.

Upon entering the host, infecting *E. coli* counteract host-mediated iron ion sequestration by secreting specialized iron chelators called siderophores, which have been detected in urine from UTI patients^11,15^. Siderophores are small molecules with affinities for iron (III) that make them competitive with mammalian iron-binding proteins^16^. Iron (III)-siderophore complexes are avidly imported by dedicated bacterial import systems to support nutritional demands for this critical, growth-limiting nutrient. Intracellular iron released from siderophore complexes then participates in feedback repression of siderophore biosynthetic genes through the ferric uptake regulator (Fur)^17,18^. The stereotypical, 19 base sequence motif (the Fur box) involved in this regulatory arrangement is a common criterion for identifying siderophore operons in genomic data^19^.

*E. coli*, as well as many other *Enterobacterales* species, secrete the catechol-based siderophore enterobactin (Ent), a siderophore with extraordinarily high iron (III) affinity (K_a_ ~10^52^)^20^. Nevertheless, *E. coli* isolates from patients with UTI encode up to three additional, genetically non-conserved siderophore systems that import siderophores yersiniabactin (Ybt), salmochelin, and aerobactin, respectively. Ybt is the non-Ent siderophore most frequently encountered in clinical urinary isolates^10,21–23^, where it is associated with increased pathogenic potential^4,24,25^. Why *E. coli* strains with elevated pathogenic potential incur the additional metabolic cost of synthesizing additional siderophores is a longstanding question. One rationale is that these non-conserved siderophores more effectively evade innate immune responses than the Ent system. Another rationale, independent of iron acquisition, is that non-conserved siderophore systems may benefit infection-associated *E. coli* strains through interactions with non-iron transition metal ions, including copper^23^ and nickel^22^. Given that a single siderophore may perform multiple functions, these rationales are not mutually exclusive.

In the present study, we ask whether Ybt is functionally redundant with Ent for iron uptake. In these experiments we compared growth of model uropathogenic *E. coli* (UPEC) strain UTI89 mutants expressing only Ent or Ybt in the presence of human transferrin. We also chemically complemented cultures of siderophore-deficient strains with purified Ent or Ybt. We used transcriptional reporter constructs for siderophore biosynthetic genes to identify a non-canonical mode of regulation consistent characteristic of quorum sensing systems. The results are consistent with a multifunctional role for Ybt that includes autoinduction in a quorum-sensing regulation in addition to transition metal ion binding.

## RESULTS

### Siderophore-dependent UPEC growth in human transferrin-containing medium

To determine whether the iron acquisition functions of UPEC siderophore systems are functionally redundant, we first sought to establish a culture condition in which growth is dependent upon siderophore production. The model UPEC strain UTI89 can secrete catecholate siderophores (enterobactin and salmochelin) and a phenolate siderophore yersiniabactin (Ybt)^24^. Enterobactin produced by UTI89 may be secreted directly or modified by C-glycosidation machinery encoded by the *iroA* cassette to form salmochelin and other modified enterobactins. To identify a siderophore-dependent growth condition we compared growth of UTI89 with its isogenic, siderophore-deficient mutant UTI89*ΔentBΔybtS*^24^. In M63/nicotinic acid/0.2% glycerol medium, which has been previously demonstrated to stimulate UPEC siderophore production^22,23,26^, UTI89*ΔentBΔybtS* exhibited only a mild growth defect in late stationary phase relative to UTI89 (see **Fig. S1a** in the supplemental material). In contrast, human transferrin (hTf) addition, previously used to create a siderophore-dependent growth condition^27^, led to a marked, highly significant UTI89*ΔentBΔybtS* growth defect relative to UTI89 (p<0.0001 at 10 hours) (**Fig. S1b**). This growth defect was not observed when hTf was replaced with heat-denatured hTf or an equivalent molar amount of bovine serum albumin (**Fig. S1c, d**), consistent with a determinative role for intact hTf iron-binding domains. FeCl_3_ supplementation (10 µM) also abolished the hTf-associated UTI89*ΔentBΔybtS* growth defect, consistent with an iron-dependent phenotype (**Fig. S1e**). Finally, substitution of hTf with 150 µM of the iron(III) chelator compound, 2,2’-dipyridyl replicated the UTI89*ΔentBΔybtS*^24^ growth defect observed with hTf (p< 0.05 at 10 hours)(**Fig. S1f**), which is also consistent with an iron-dependent phenotype. Together, these data are consistent with M63/nicotinic acid/0.2% glycerol/hTf (hereafter abbreviated as M63+hTf) as a siderophore-dependent growth condition.

### Siderophore system contributions to siderophore-dependent growth

To determine whether the Ent and Ybt systems are similarly capable of supporting siderophore-dependent growth, we compared growth between UTI89 mutants expressing Ent or Ybt as the sole siderophore (UTI89*ΔiroAΔybtS* and UTI89*ΔentB*, respectively) (**Table 1**). In the absence of hTf, both mutants grew similarly to UTI89 (**Fig. 1a**). In the presence of hTf, UTI89*ΔiroAΔybtS* grew similarly to UTI89, but UTI89*ΔentB* exhibited a profound growth deficiency comparable to the siderophore-deficient strain UTI89*ΔentBΔybtS* (p<0.0001 at 10 hours) (**Fig. 1b**). Comparable results were obtained with substitution of 2,2’-dipyridyl for hTf (**Fig. S2**). Together, these results are inconsistent with the hypothesis that the Ybt siderophore system is functionally redundant with Ent system-mediated iron acquisition. The results are consistent with a role for Ent, but not Ybt, in supporting UTI89 growth from low cellular density in the presence of hTf.

**Table 1:**
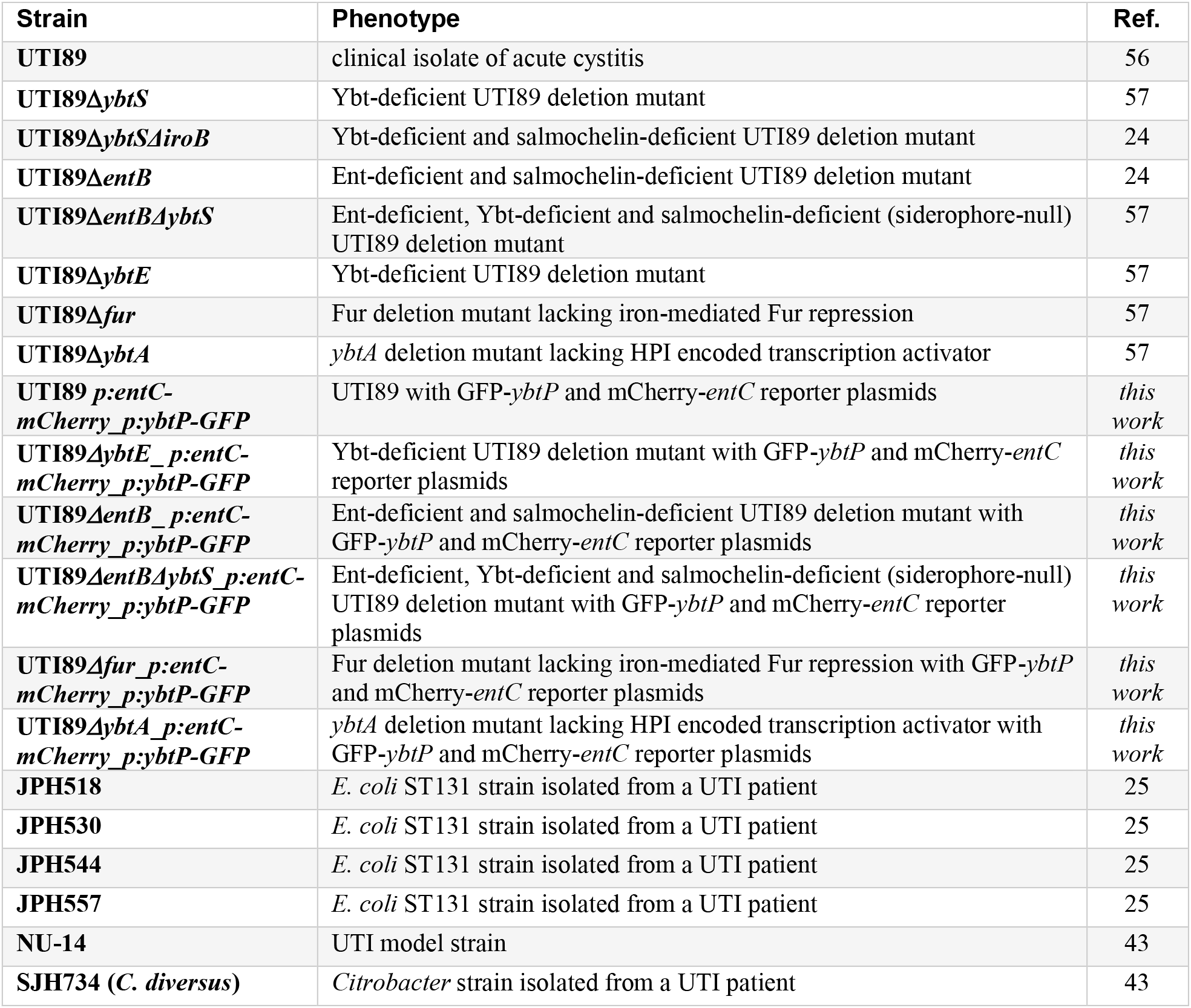
Bacterial strains used in this study.

**Figure 1.**
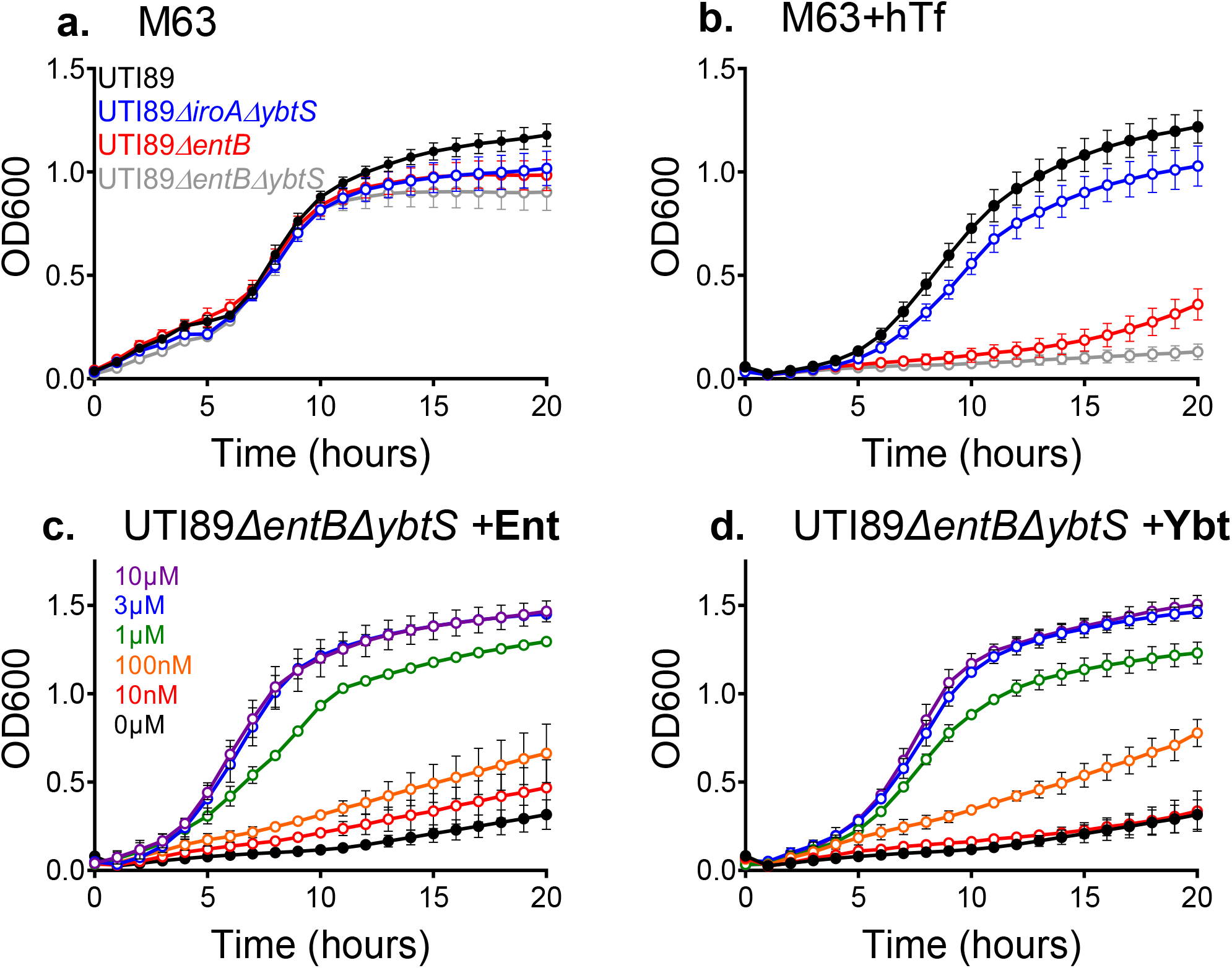
Yersiniabactin (Ybt) biosynthesis does not compensate for loss of enterobactin (Ent) biosynthesis during siderophore-dependent growth of *E. coli* UTI89 and isogenic mutants. (a,b) Growth curves of UTI89 and isogenic siderophore biosynthesis mutants in M63/glycerol media in the absence (a) or presence of 3 μM human *apo*-transferrin (hTf, b). (c,d). Growth curves of the complete siderophore biosynthesis-deficient mutant UTI89*ΔentBΔybtS* in M63+hTf containing increasing concentrations of purified Ent (c), or Ybt(d).

### Chemical complementation effects on siderophore-dependent growth

We considered that non-redundancy of the Ybt system may arise from deficient Ybt siderophore activity relative to Ent. To test this hypothesis, we chemically complemented UTI89*ΔentBΔybtS* in M63+hTf with equimolar quantities of either purified Ent or Ybt. We observed a comparable dose-response relationship for the two siderophores, reaching a maximal response at 3 μM for each (**Fig. 1c,d**). This result is consistent with Ybt and Ent having equivalent siderophore activity in M63+hTf. Thus, the growth defect of UTI89*ΔentB* in M63+hTf (**Fig. 1b**) is not due to an inability of Ybt to substitute for Ent’s siderophore activity.

### Extracellular siderophore accumulation during siderophore-dependent growth

We next considered the possibility that Ent and Ybt production differ in our siderophore-dependent culture conditions. To compare siderophore production in M63+hTf, we used liquid chromatography-mass spectrometry to quantify siderophores in conditioned media over time. In UTI89 cultures, Ybt concentrations exhibited a delayed increase compared to Ent, paralleled the growth (optical density) curve, and were maximal during stationary phase (**Fig. 2a**). In the Ent-deficient mutant UTI89*ΔentB*, Ybt production kinetics were markedly depressed (**Fig. 2b**). Conversely, Ent production kinetics were substantially unchanged in the Ybt-deficient mutant UTI89*ΔybtSΔiroA* (**Fig. 2c**). In the absence of hTf, Ybt production kinetics by UTI89*ΔentB* and Ent production kinetics by UTI89*ΔybtSΔiroA* were both comparable to UTI89 (**Fig. S3a,b,c**). These results are consistent with Ybt production during low bacterial cell density culture that is insufficient to overcome the absence of Ent during siderophore-dependent growth.

**Figure 2.**
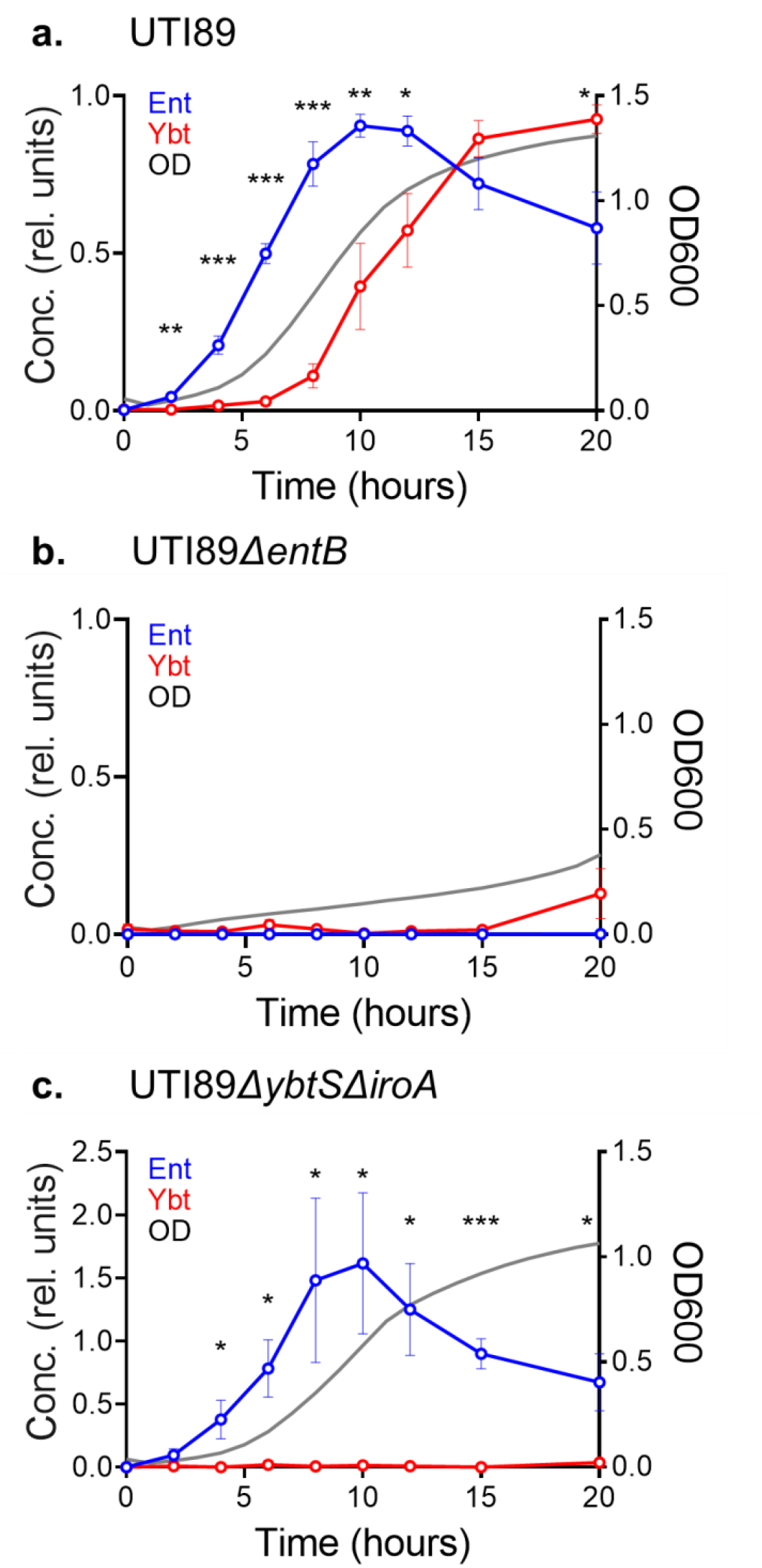
Time course of Ent and Ybt production by *E. coli* UTI89 and isogenic mutants. Liquid chromatography-mass spectrometry was used to quantify Ent (blue) and Ybt (red) content of M63+hTf medium conditioned by wild type UTI89 (a), UTI89*ΔentB*(b), or UTI89*ΔybtSΔiroA*(c) over 20 hours. Concentration values are relative to maximum values in UTI89. Culture density (optical density at 600 nm) is depicted by the gray plot line. Statistical comparison between relative conc. of Ent and Ybt at each time point: *, *P* < 0.05; **, *P* < 0.01; ***, *P* < 0.001 by unpaired T-test.

### Ybt biosynthetic gene transcription is delayed during siderophore-dependent growth

The disparate growth kinetics displayed by UTI89 siderophore mutants may reflect differences in siderophore biosynthetic gene expression during early culture. To test early biosynthesis gene expression, we used qRT-PCR to compare mRNA transcripts from early Ent and Ybt biosynthesis genes (*entB* and *ybtS*, respectively) (**Table S1**). To accommodate the large media volumes necessary to obtain adequate RNA yield from low density cultures, we substituted hTf with 2,2’-dipyridyl. At one hour of culture, the expression fold change of *entB* was significantly higher than that of *ybtS* (p=0.03). By four hours, this difference was diminished and became insignificant (p>0.05) (**Fig. 3**). This finding is consistent with a UTI89 siderophore system transcriptional response that emphasizes Ent biosynthesis during early growth and Ybt biosynthesis later in culture when cell density is greater.

**Figure 3.**
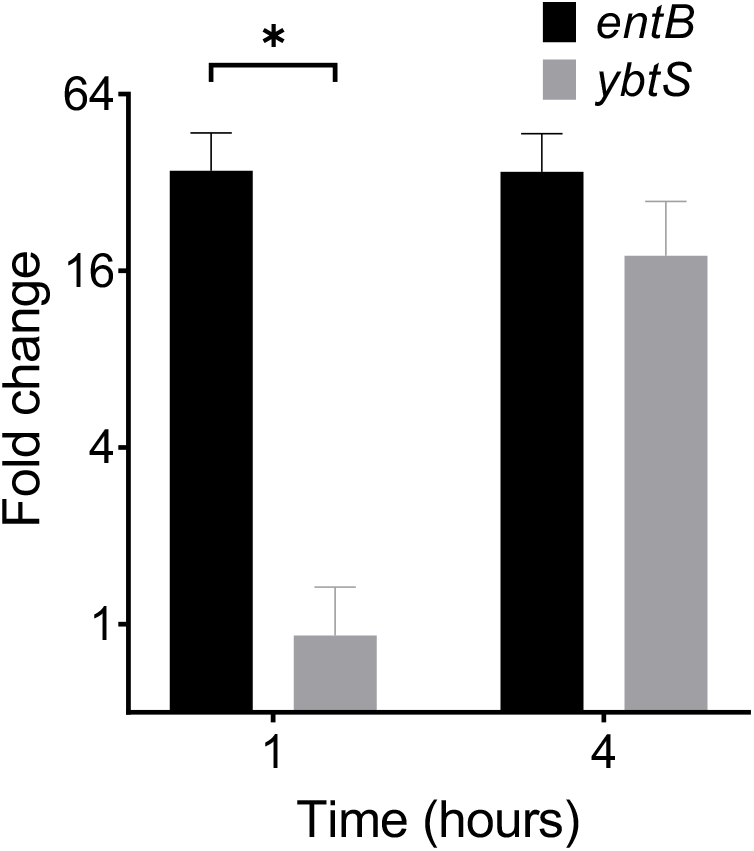
Early siderophore biosynthetic gene transcription in UTI89 assessed by qRT-PCR during siderophore-dependent growth. mRNA fold change of *entB* and *ybtS* biosynthesis gene expression at 1 and 4 hours in M63 media containing 150 µM 2,2’-dipyridyl, normalized to 10 hour time point. *rssA* was used as a housekeeper gene. *, *P* < 0.05 using unpaired T-test.

### Ybt biosynthetic gene upregulation delay is typical of culture populations

To determine whether Ybt expression was delayed uniformly by cells in culture or is attributable to distinct subpopulations, we created reporter plasmids in which Ent or Ybt biosynthetic operon promoters drive expression of different fluorescent reporters and measured their activity with flow cytometry (**Fig. S4a,b**) (**Table S1**). In *p:entC-mCherry*, the *entCEBA* promoter for the main Ent biosynthetic operon controls mCherry expression. In *p:ybtP-GFP* the *Yersinia* HPI operon 1 promoter controlling transcription of the first Ybt biosynthetic gene (*ybtS*) controls GFP expression^28^(**Fig. 4a-d**). In M63+hTf, mean fluorescence intensity (scaled to maximum value) of the *Yersinia* operon 1 reporter in UTI89 *p:entC-mCherry p:ybtP-GFP* was significantly delayed over the first 8 hours compared to the *entCEBA* reporter (2 hours p=0.054, 4-8 hours p<0.05, 10 hours p=0.076) (**Fig. 4e)**. These results are consistent with the qRT-PCR results and demonstrate a population-wide delay in Ybt biosynthetic gene upregulation by UTI89 in M63+hTf. Differences in transcriptional activation may relate to inability of the Ybt system to compensate for the loss of Ent during siderophore-dependent growth.

**Figure 4.**
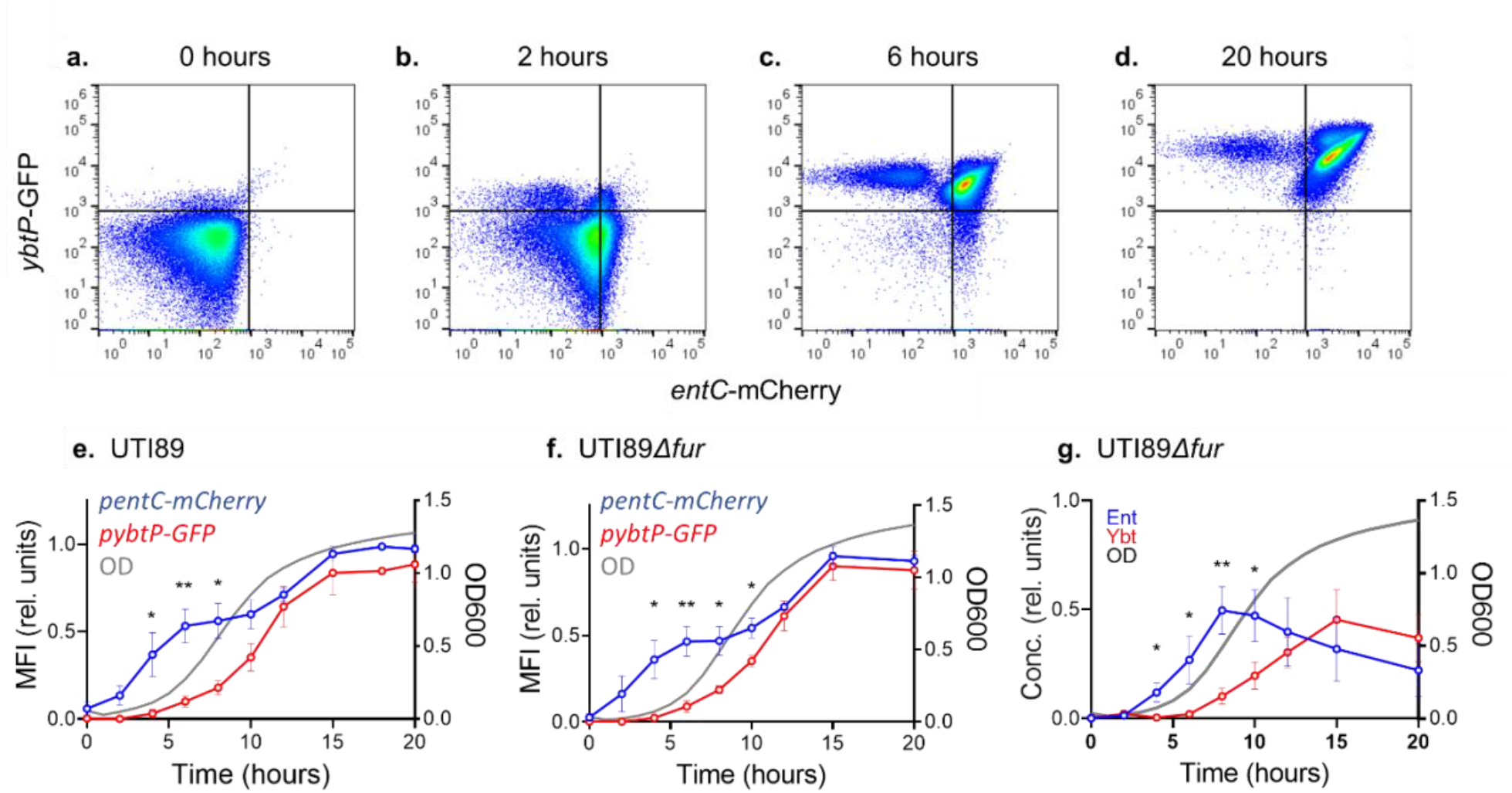
Siderophore biosynthetic operon transcription in UTI89 and its *fur*-deficient mutant assessed by fluorescent reporter expression during siderophore-dependent growth. Fluorescent protein expression in bacterial cultures was measured using flow cytometry. GFP fluorescence controlled by the *Yersinia* operon 1 promoter (*ybtP*-GFP) and mCherry fluorescence controlled by the *entCEBA* promoter (*entC-*mCherry) are displayed. (a-d) Representative pseudocolor plots of UTI89 *p:entC-mCherry p:ybtP-GFP* at 0, 2, 6, and 20 hours. Quadrants are gated on non-fluorescent protein cultures. (e,f) Mean fluorescence intensity (MFI) for each reporter of normalized to WT during growth of UTI89 *p:entC-mCherry p:ybtP-GFP* (e) or UTI89*Δfur p:entC-mCherry p:ybtP-GFP* (f) in M63+hTF. (g) Ent and Ybt concentrations relative to wild type (UTI89) in M63+hTf medium conditioned by UTI89*Δfur*. Culture density (optical density at 600 nm) is depicted by the gray plot line. Statistical comparison between relative MFI of *p:entC-mCherry* and *p:ybtP-GFP* at each time point: *, *P* < 0.05; **, *P* < 0.01; ***, *P* < 0.001 by unpaired T-test.

### Fur repression does not account for delayed ybt production and expression

We next examined how transcriptional control of Ent and Ybt biosynthesis differs. Fur (ferric uptake regulator) is the canonical siderophore system regulator and participates in homeostatic feedback repression by iron, though more complex regulatory schemes are possible^29–31^. To determine whether more profound Fur repression explains delayed Ybt system transcription, we compared reporter expression and siderophore production between UTI89 and its Fur-deficient mutant UTI89*Δfur*. We observed that the late *Yersinia* operon 1 reporter stimulation observed in UTI89 persists in UTI89*Δfur* when compared to *entCEBA* reporter (4-10 hours p<0.05) (**Fig. 4e**,**f**). There remains a corresponding, significant delay in Ybt production relatively to Ent (4-10 hours p<0.05) (**Fig. 4g**). The retained delay in Ybt biosynthetic upregulation is inconsistent with differential Fur regulation as the basis for kinetic differences in Ent and Ybt biosynthesis in these conditions.

### Ybt biosynthetic gene upregulation by Ent-null mutant is abolished during siderophore-dependent growth

To determine whether transcriptional Ybt biosynthetic upregulation occurs in the UTI89*ΔentB* background, we monitored reporter activity in UTI89*ΔentB p:entC-mCherry p:ybtP-GFP* during growth in M63+hTf. In this strain, the increase in *Yersinia* operon 1 reporter MFI observed with the wild type reporter strain was abolished for the duration of the assay (p<0.05, 10-20 hrs) (**Fig. 5a**). This is notable as there is no known direct regulatory connection between the two siderophore systems and Ent-null UTI89 can grow and produce Ybt comparably to wild type UTI89 in the absence of competitive iron chelators (**Fig S3**) ^24^. We considered an indirect connection between the two siderophore systems in the M63+hTf condition. We noted that differences in Ybt biosynthetic transcriptional activity between the wild type and Ent-deficient reporter strains coincide with differences in culture density (5 hours p=.02, 6 hours p=.002, 7-20 hours p<0.001)(**Fig. 5b**). Ent stimulation in the Ent deficient mutant still expressed *entCEBA* reporter at similar level during growth (0-15 hours p>0.05) **(Fig. S5)**. This suggested to us the possibility of a density-dependent regulatory input on Ybt biosynthesis.

**Figure 5.**
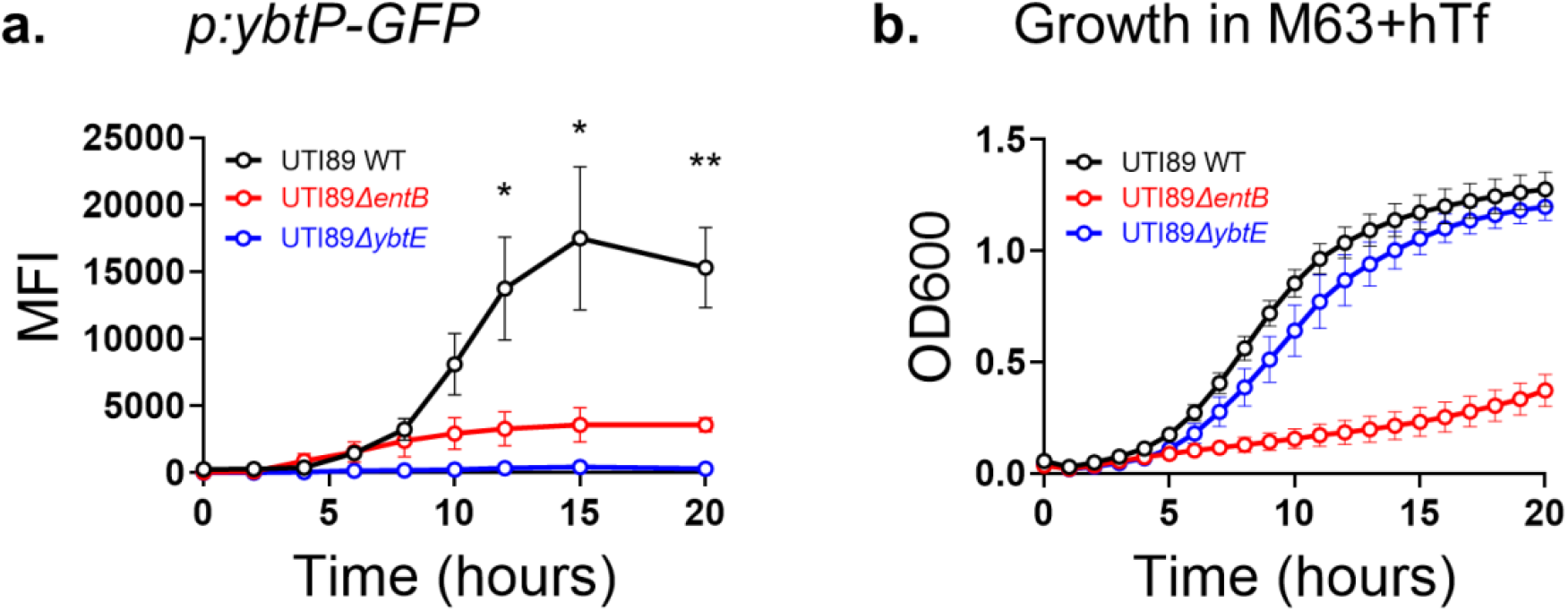
Lack of growth or loss of Ybt biosynthesis represses *Yersinia* operon 1 expression. UTI89 strains carrying the *p:entC-mCherry* and *p:ybtP-GFP* plasmids were grown in M63+hTf. *p:ybtP-GFP* reporter expression is displayed as mean fluorescence intensity (MFI) of flow cytometry measurements (a) and growth measured by optical density (OD600, b). Statistical comparison between WT and both mutant strains of the MFI of *p:ybtP-GFP* at each time point: *, *P* < 0.05; **, *P* < 0.01; ***, *P* < 0.001 by unpaired T-test.

### Ybt is an autoinducer

We hypothesized that an important Ybt biosynthetic stimulus derives from the density-dependent regulation characteristic of a quorum sensing (QS) system. A defining feature of QS systems is production of a secreted molecule that accumulates outside the cell in proportion to population density. This secreted molecule – an autoinducer - then upregulates its own production in a concentration-dependent manner, creating a feed-forward autoregulatory loop. Although multiple QS systems have been described in *E. coli*^32–35^, we hypothesized that Ybt fulfills the autoinducer role in addition to its siderophore role. To test this, we compared *Yersinia* operon 1 reporter activity between UTI89 *p:entC-mCherry p:ybtP-GFP* and the Ybt biosynthesis-null mutant UTI89*ΔybtE p:entC-mCherry p:ybtP-GFP* during siderophore-dependent growth in M63+hTf. Despite similar growth curves, the GFP reporter was markedly suppressed in UTI89*ΔybtE* (**Fig. 5a)**. At 20 hours MFI of *Yersinia* operon 1 reporter was significantly diminished compared to UTI89 WT **(Fig. 6a,d)**. Chemical complementation of UTI89*ΔybtE p:entC-mCherry p:ybtP-GFP* with purified Ybt restored GFP fluorescence (**Fig. 6b,d**), while equimolar Ent did not (**Fig. 6c,d**). Together, these results distinguish Ybt production as subject to autoinductive regulation in which Ybt plays a dual role as effector and signaling molecule. In this context, deficient Ybt accumulation in the medium leads to inadequate Ybt autoinduction at low population density. Ent production, in contrast, is activated independently of population density, facilitates entry into logarithmic growth in M63+hTf, and facilitates higher Ybt production by the larger bacterial population.

**Figure 6.**
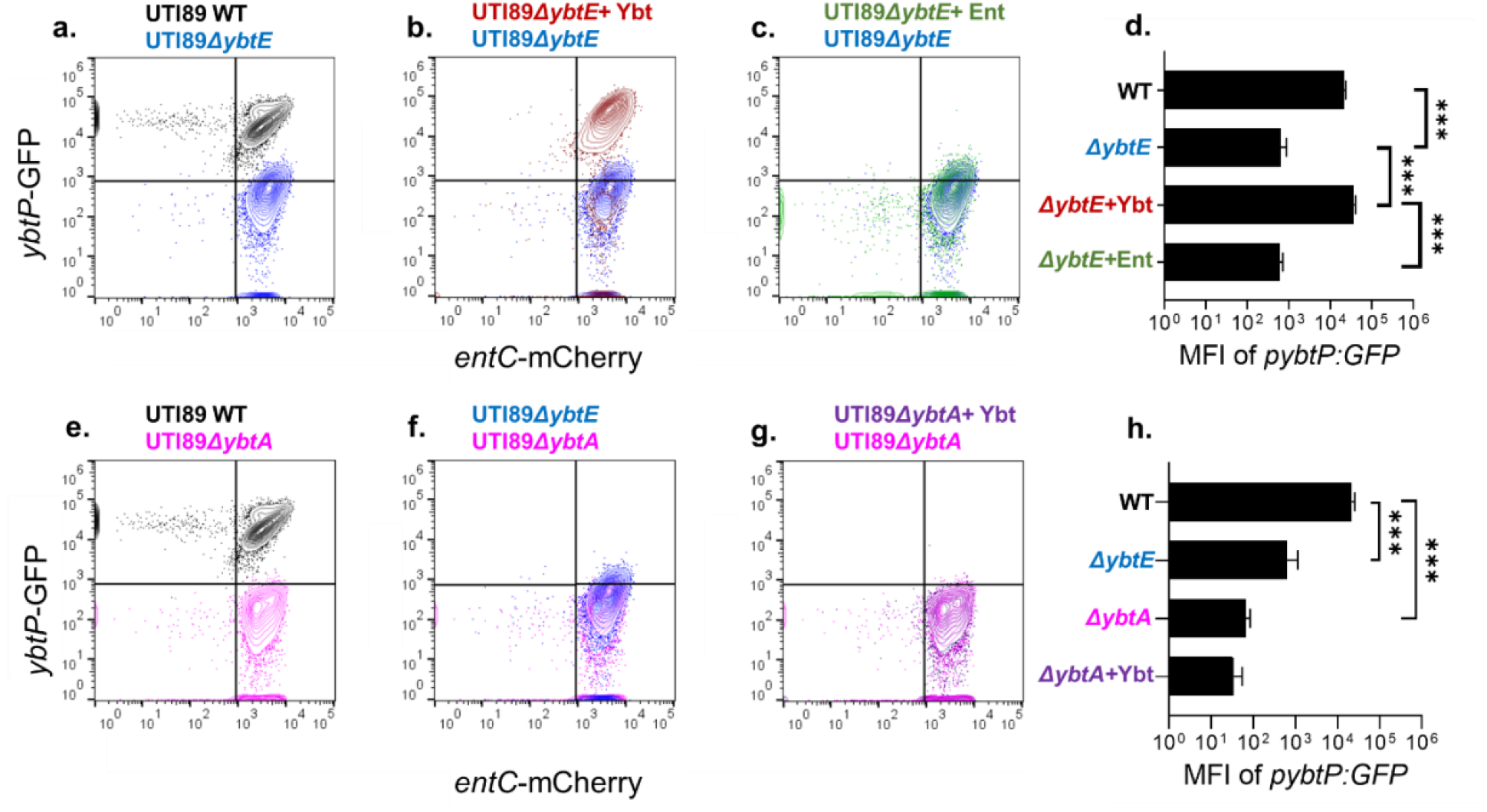
Siderophore biosynthetic operon transcription in UTI89, UTI89 mutants, and with chemically complemented cultures assessed by fluorescent reporter expression. (a-c, e-g) Representative pseudocolor plots of strains transfected with *p:entC-mCherry* and *p:ybtP-GFP* after 20 hours of growth in M63+hTf. Quadrants are gated on non-fluorescent protein cultures. (d,h) Mean fluorescence intensity (MFI) of *ybtP-*mCherry from each strain at 20 hours. Culture medium was supplemented with 10 nM Ybt (+Ybt, b,d,g,h) or Ent (+Ent, c,d). *, *P* < 0.05; **, *P* < 0.01; ***, *P* < 0.001 by unpaired T-test.

### YbtA is a candidate Ybt sensor

Most QS systems follow a general functional model for density-dependent regulation^36–39^. Applying this paradigm to the Ybt system, we anticipated a Ybt-specific response element that increases Ybt biosynthesis. A prominent candidate for this receptor is the AraC-type transcription factor YbtA, which is predicted to possess a ligand binding domain that may act as a Ybt sensor^28,40,41^. To assess the role of YbtA in controlling Ybt biosynthesis, we measured reporter activity in the YbtA-deficient mutant UTI89Δ*ybtA*. Compared to UTI89, *Yersinia* operon 1 reporter activity in UTI89Δ*ybtA* remained minimal throughout M63+hTf cultures (**Fig. 6e,h**), with a non-significant trend toward lower reporter activity than UTI89Δ*ybtE* at 20 hours (p=0.12)(**Fig. 6f,h)**. Unlike UTI89Δ*ybtE*, the loss of *Yersinia* operon 1 reporter activity in UTI89Δ*ybtA* was not restored by addition of purified Ybt to the culture medium (**Fig. 6g,h**).

Although these results do not definitively identify YbtA as a Ybt receptor, they are consistent with this possible role and show that YbtA is necessary for Ybt biosynthetic gene transcription.

### Differential Ent and Ybt production is typical of multiple clinical urinary isolates

To determine whether other urinary isolates carrying the *Yersinia* HPI exhibit density-dependent Ybt biosynthetic regulation similar to UTI89, we measured siderophore production by genetically distinctive clinical urinary isolates cultured in M63+hTf (Table 1)^42,43^. In addition to *E. coli*, this included *Citrobacter diversus*, a distinctive *Enterobacterales* species (**Fig. 7f**). We compared siderophore concentrations of Ent and Ybt during early (8 hours) and late (15 hours) time points. As with UTI89, the Ybt:Ent ratio was higher in late than in early time points for all strains examined (**Fig. 7a-f**). These findings are consistent with density-dependent Ybt production in multiple urinary *Enterobacterales* isolates, similar to that observed in UTI89.

**Figure 7.**
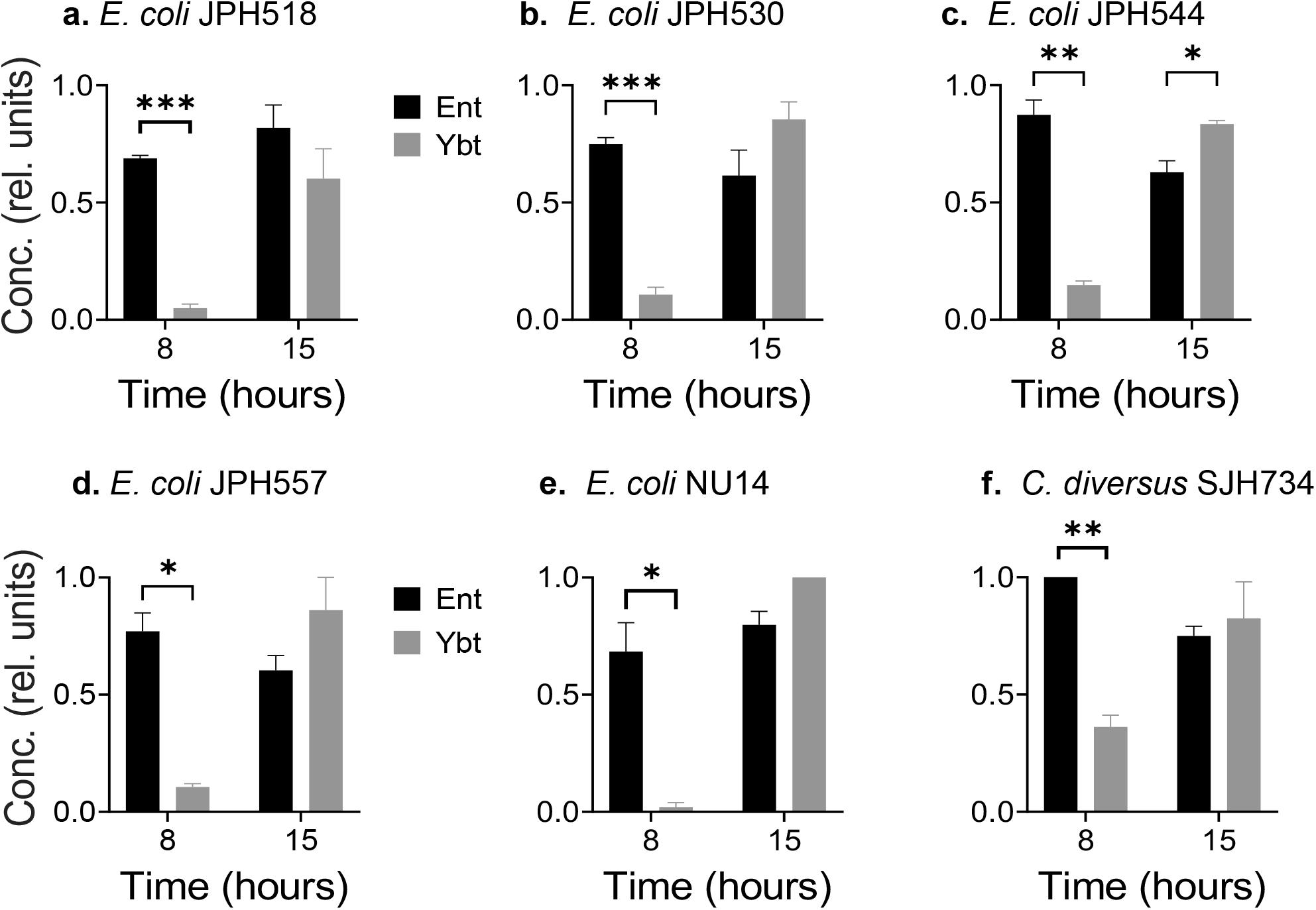
Siderophore production by clinical urinary isolates. Ent and Ybt concentrations relative to UTI89 in M63+hTf medium. Results at early (8 hour) and late (15 hour) time points are displayed for different *E*.*coli* (a-e) and *Citrobacter diversus*, f). *, *P* < 0.05; **, *P* < 0.01; ***, *P* < 0.001 by unpaired T-test.

## DISCUSSION

We find that, during siderophore-dependent growth of UPEC, the Ybt system is not fully functionally redundant with the genetically conserved Ent system. This is attributable to deficient Ybt biosynthesis at low cell density. Full Ybt biosynthesis instead requires a density-dependent transcriptional regulatory cycle in which Ybt acts as an autoinducer (**Fig. 8**). These characteristics are consistent with the quorum-sensing regulation, where cellular crowding leads to extracellular accumulation of an autoinductive signal. While QS control of siderophore system activity has been described in other bacteria^36,44–46^, this is the first description of QS signaling and siderophore activity encoded by the same genetic unit, using the same extracellular small molecule effector.

**Figure 8:**
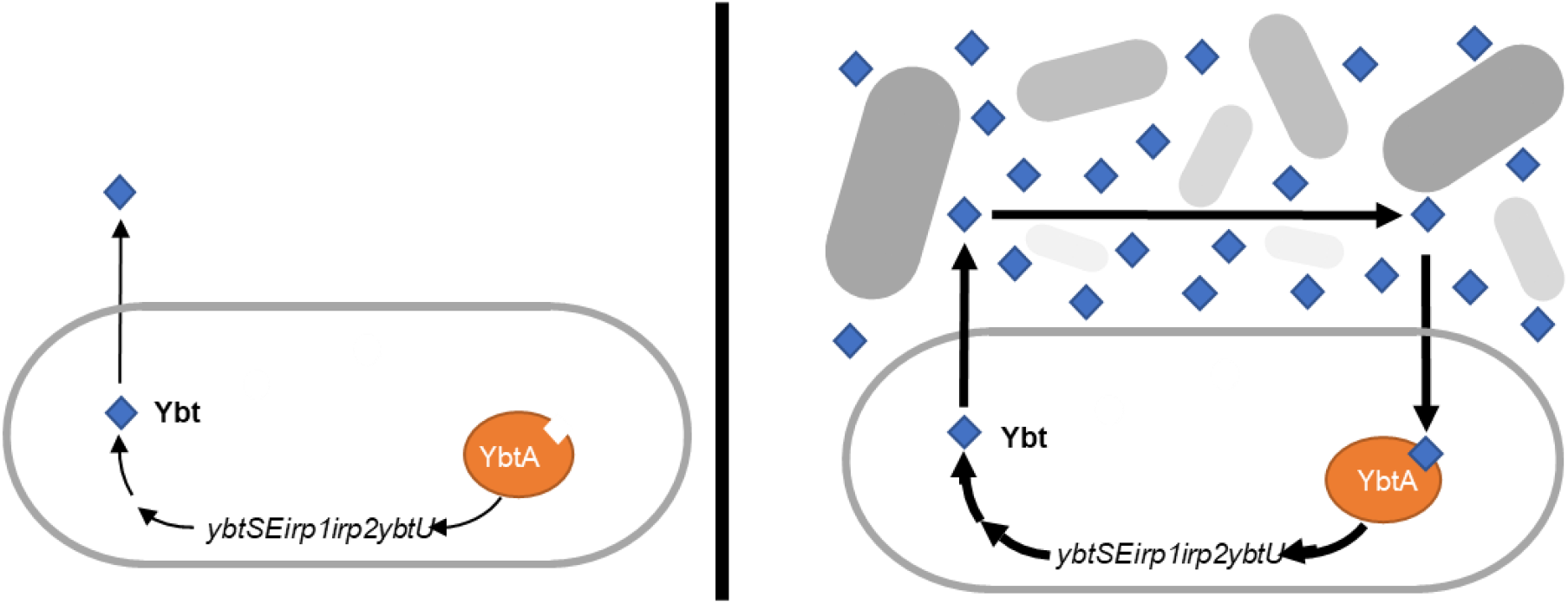
Model of Ybt-mediated quorum sensing autoregulation. At low cell density and limited iron availability (*left*), Ybt biosynthesis is insufficient to match the iron uptake activity of the Ent system. Extracellular Ybt accumulates slowly and autoinduction is minimal. With increasing cell density (*right*), extracellular Ybt accumulates sufficiently to bind YbtA and stimulate increased Ybt biosynthesis by each cell. This autoinductive cycle rapidly increases extracellular Ybt as cell density increases.

A combined siderophore and QS system may possess distinctive advantages. Encoding both functions within a single non-conserved, mobile genetic island is genetically efficient, facilitating horizontal gene transfer. In the canonical model involving a freely diffusible siderophore, this arrangement also limits the metabolic cost of Ybt biosynthesis in conditions where Ybt diffuses away and is unlikely to retrieve or control metal ions in the immediate environment. In higher cell density environments, Ybt sharing among producers and recycling of imported Ybt permits more rapid autoinducer accumulation while minimizing biosynthetic costs^23^. This view of Ybt as a “public good” contrasts with the Ent system, which *E. coli* appear capable of using as a “private good” at low cell density^47^ and requires Ent hydrolysis to release imported iron. Density-dependent siderophore activity may be of particular value in iron-limited host niches where bacterial crowding or confinement is typical. These environments may include epithelial surfaces, biofilms, and intracellular compartments. Consistent with this scenario is the previous finding that *ybtS* is the most upregulated UTI89 gene in biofilm-like intracellular bacterial communities (IBCs) residing within bladder epithelial cells during experimental murine UTI^48^.

QS stimulation may also optimize Ybt interactions with non-iron transition metal ions such as copper. This may be advantageous during bacterial compartmentalization in the phagolysosome where macrophage-like cells intoxicate bacteria with copper, making Ybt chelation especially advantageous. UPEC compartmentalization may thus serve as an environmental cue, engaging the Ybt autoinductive response and maximizing intraphagosomal copper-Ybt (Cu-Ybt) complex formation. In addition to controlling copper disposition, Cu-Ybt catalyzes superoxide dismutation similarly to superoxide dismutases^49^. Cu-Ybt has been recently described to stimulate Ybt biosynthesis^28^, possibly through YbtA-dependent Cu-Ybt stimulation, augmented autoinduction, and/or signaling by copper ions released from the complex^23,26^. Whether the Ybt system persists among UPEC due to its iron scavenging activity, its interactions with non-iron metal ions, or both, remains unclear.

QS system regulation of siderophore systems has been previously described in other bacterial orders, where the autoinducer and the siderophore are different molecules. In the prototypical quorum-sensing bacterium *Vibrio harveyi*, siderophore production is regulated by the canonical Lux QS system^44^ which, unlike the Ybt system, represses siderophore expression with increasing cell density. In *Pseudomonas*, the related LasR QS system increases production of the siderophore pyoverdine with increasing cell density^45^. Both the Lux and Las systems use acylhomoserine lactones as the autoinducer. These varying responses to cellular density may reflect adaptation to different environmental stresses and possibly coordination with different, alternative iron acquisition strategies. Nevertheless, the chemical and regulatory diversity of siderophore systems suggest that bacteria have evolved many strategies to control, acquire, and use transition metal ions in their environments.

The *Yersinia* HPI, which is more complex than many siderophore systems, encodes components that are typical of QS system, specifically autoinducer biosynthesis, autoinducer transport, and intracellular receptor/regulator functions^50^. The precise functions of these components and their interactions with non-HPI components are incompletely understood. Previous studies have confirmed that Ybt is internalized by an outer membrane transporter (FyuA)^23,51^, suggesting the existence of an intracellular receptor. The most likely candidate for this is YbtA, an AraC-type transcription factor with a well-conserved DNA binding domain, which is encoded as an independent operon (operon 2) in the HPI and controls transcription of multiple HPI operons^40,52,53^. Consistent with the autoregulatory function of a QS system, YbtA is predicted to possess an N-terminal ligand-binding domain typical of AraC family proteins that is a strong candidate Ybt receptor candidate^18,28,40^. We speculate that binding to this domain of Ybt, or a derivative thereof, increases transcription of HPI operons and increases Ybt biosynthesis. Further investigation is necessary to test this hypothesis and to construct a more detailed model for its function in *E. coli* and related *Enterobacterales*, which will help us to better understand the host factors exerting selective pressure on these bacteria.

## MATERIALS AND METHODS

### Bacterial strains, and culture conditions

Bacterial strains, including UTI89 and previously characterized deletion mutants, used in this study are listed in Table 1. For vector transformations, starter cultures were grown on LB agar with antibiotics as appropriate overnight at 37 °C. Ampicillin (100 µg/ml; Gold Biotechnology), chloramphenicol (34 µg/ml; Gold Biotechnology), and/or kanamycin (100 µg/ml; Gold Biotechnology) we used for selection.

### Bacterial growth curves

Bacteria cultures (3 ml LB medium) were grown overnight with continuous shaking at 37 °C. Cells were collected from 1 ml of culture and resuspended in 3 ml of M63 minimal media (0.5 M potassium phosphate, pH 7.4, 10g/liter (NH4)2SO4, 2 mM MgSO4, 0.1 mM CaCl2, 0.2% glycerol, 10 g/ml niacin)^24^. The cells were then grown for 4 hours shaking in 37 °C. 1 ml of cells were collected and washed with fresh M63 media. Before adding the cells to the 96-well round bottom plate, the wells were filled with 200 µl fresh M63 media with or without 3 µM of human transferrin (Sigma-Aldrich #T1147) and incubated at room temperature for 30 minutes. Bacteria were added to the 96-well plate for a starting concentration 0.01 OD (around 8 million CFU). The plate was then grown shaking at 37 °C for 20 hours with hourly A_600_ OD measurements using a Tecan SPARK multimode microplate reader with a modular design including an incubator, shaker, and lid lifter (catalog #30086376).

### Enterobactin extraction

Enterobactin was isolated using method described in previous publications^10,54^. Fractions containing pure Ent were collected, lyophilized, and resuspended in water. Concentrations of metal-free Ent were determined using the Chrome Azurol S assay^55^.

### Yersiniabactin extraction

Yersiniabactin was isolated using method describe in pervious publication^28^. Fractions containing pure Ybt were collected, lyophilized, and resuspended in water. Concentrations of metal-free Ybt were determined using the Chrome Azurol S assay^55^.

### Liquid chromatography-mass spectrometry (LC-MS)

LC-MS analyses was conducted with an HPLC–equipped AB Sciex 4000 QTRAP with a Turbo V ESI ion source run in positive ion mode (Shimadzu, Kyoto, Japan). The samples were injected onto a phenylhexyl column (100 by 2.1 mm, 2.7 µm particle) (Ascentis Express, Supelco, Bellefonte, PA) with a flow rate of 0.4 ml/min. The following gradient was used: solvent A (0.1% [vol/vol] formic acid) was held constant at 95% and solvent B (90% [vol/vol] acetonitrile, 0.1% [vol/vol] formic acid) at 5% for 2 min. Solvent B was increased to 65% by 6 min and to 98% by 8 min. Solvent B was then held constant at 98% until 9 min before it was decreased to 5% by 11 min. Solvent B was then held constant at 5% for 1 additional minute. The collision energy was set at 37 V.

### qRT-PCR of bacterial mRNA

UTI89 was grown in 500 ml of M63 minimal media with 150 µM 2,2’-dipyridyl shaking at 37 °C. RNA was extracted from bacterial cultures using Qiagen RNA isolation kit. RNA was converted to cDNA, using a Thermo Fisher thermocycler, qRT-PCR was run on the cDNA. Primers for *ybtS, entB*, along with a housekeeper *rssA*, were used to measure siderophore biosynthesis transcription (**Table S1**).

### Fluorescent reporter constructs

Protein expression vector pMAL-c5Xa (NEB) was used as the backbone for constructing the mCherry reporter. The malE gene and the lacI promoter were restricted from the vector using SacI and KasI restriction enzymes. The *entCEBA* promoter was amplified with primers GK073-F/GK073-R (**Table S1**) and inserted into the pMAL-c5Xa backbone. The reporter constructs were transformed into respective strains as indicated. An inducible GFP was made by inserting the *Yersinia* operon 1 promoter sequence into plasmid pFCcGi (Addgene) upon restriction with HindIII and XbaI (NEB). The resulting construct was called *pybtP:GFP*^28^. The reporter constructs were transformed into respective strains as indicated.

### Flow Cytometry

The bacterial cells isolated from the bacterial cultures were resuspended in 4% paraformaldehyde PBS to fix the cells. The cells were incubated for 30 minutes at room temperature. After 30 minutes, 1 ml of cold PBS was added to quench the fixative. The cells were resuspended in cold FACS buffer (1% BSA, 0.1% sodium azide in PBS). The cells were then measured on a Cytek Aurora® flow cytometer (N9-20006 Rev. B) using violet, blue, yellow-green, red lasers. SpectroFlo® software and quality control beads were used to control and read the data from the flow cytometer. The data was then analyzed using FlowJo software. Negative FACS gates were determined using non-fluorescent protein contain cultures.

## ACKNOWLEDGEMENTS

J.P.H acknowledges National Institute of Health (NIH) grant R01DK111930 and RO1DK125860.

W.H.M. acknowledges NIH funding sources: KL2TR002346, UL1TR002345, 5K08AR076464. We acknowledge Anne Robinson and John Robinson for their insightful discussions. We acknowledge Hung Tran and Anthony Tran for helping make bacterial mutant constructs. We declare no conflicts of interest.

---

**Figure S1:**
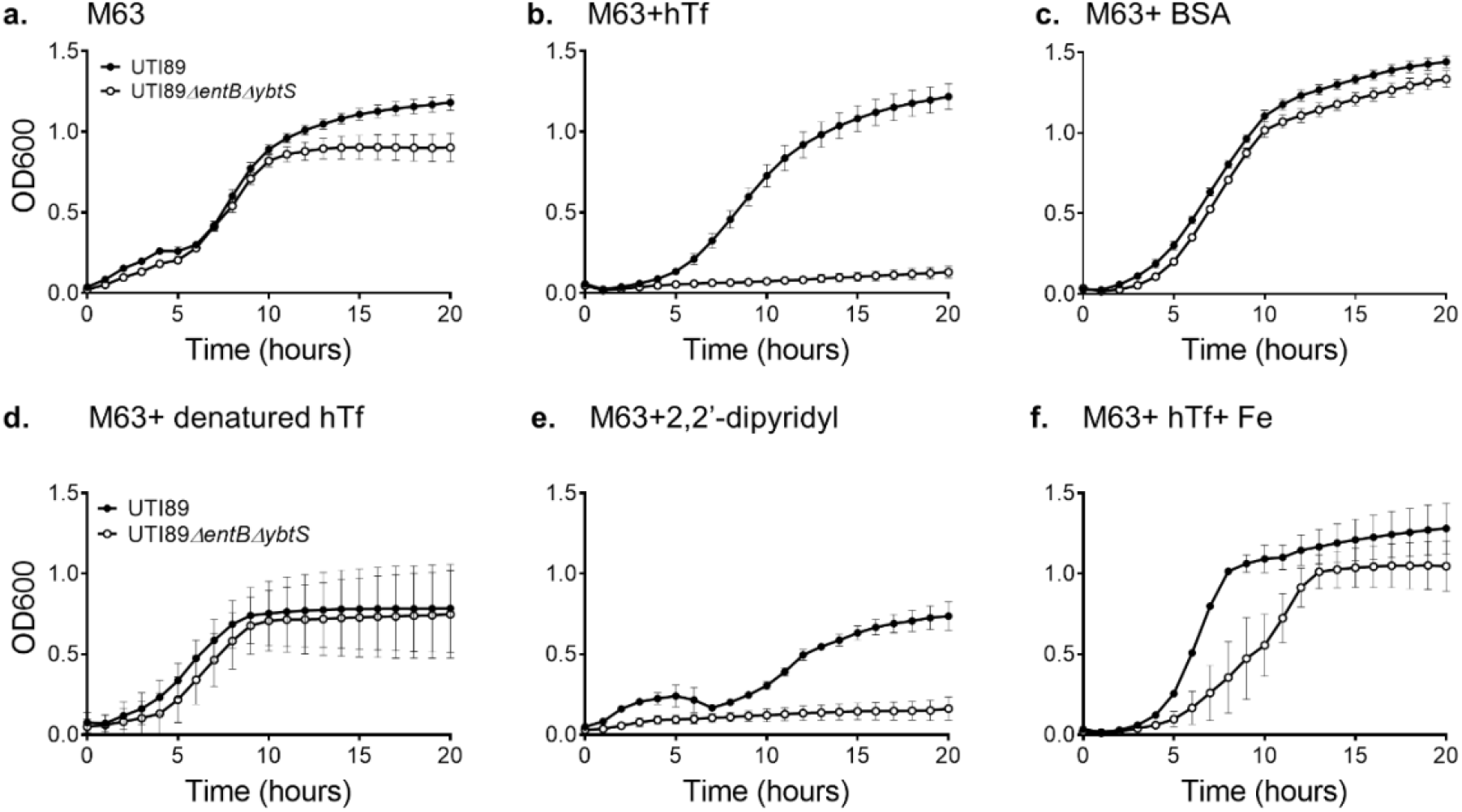
Siderophore-null UTI89 strain does not grow in media supplemented with human transferrin or the iron ion chelator 2,2’-dipyridyl. Growth curves of wild type UTI89 (black) and siderophore-null UTI89ΔentBΔybtS (white) in M63 media alone (a), with 3 μM human Transferrin (hTf) (b), 3 μM bovine serum albumin (BSA)(c), 3 μM heat-denatured hTf (d), 150 μM 2,2’-dipyridyl (e), or 3 μM hTf and 10 μM FeCl_3_(f). Cultures were grown in a 96-well plate shaking for 20 hours at 37 °C. Wells were measured at 600 nm wavelengths every hour by a Tecan plate reader.

**Figure S2:**
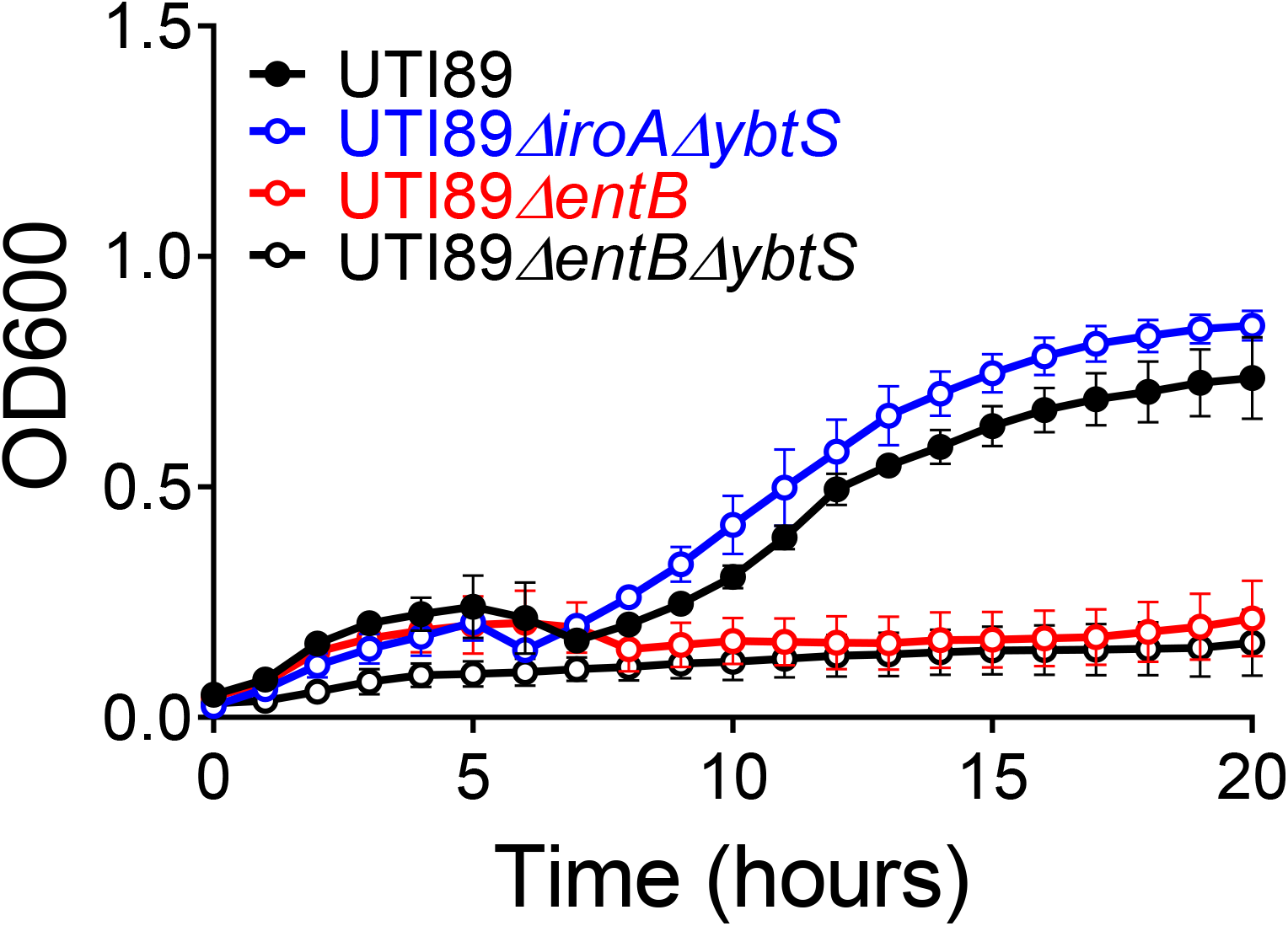
Ybt-only producing strain is diminished in iron-limiting conditions. Growth curve of UTI89 strains with potential to make Enterobactin (Ent), Salmochelin, and Yersiniabactin (Ybt) (black), Ent only (blue), Ybt only (red), or no siderophores (white). Strains were grown in M63+ 150 μM 2,2’-dipyridyl. Cultures were grown in a 96-well plate shaking for 20 hours at 37 °C. Wells were measured at 600 nm wavelengths every hour by a Tecan plate reader.

**Figure S3:**
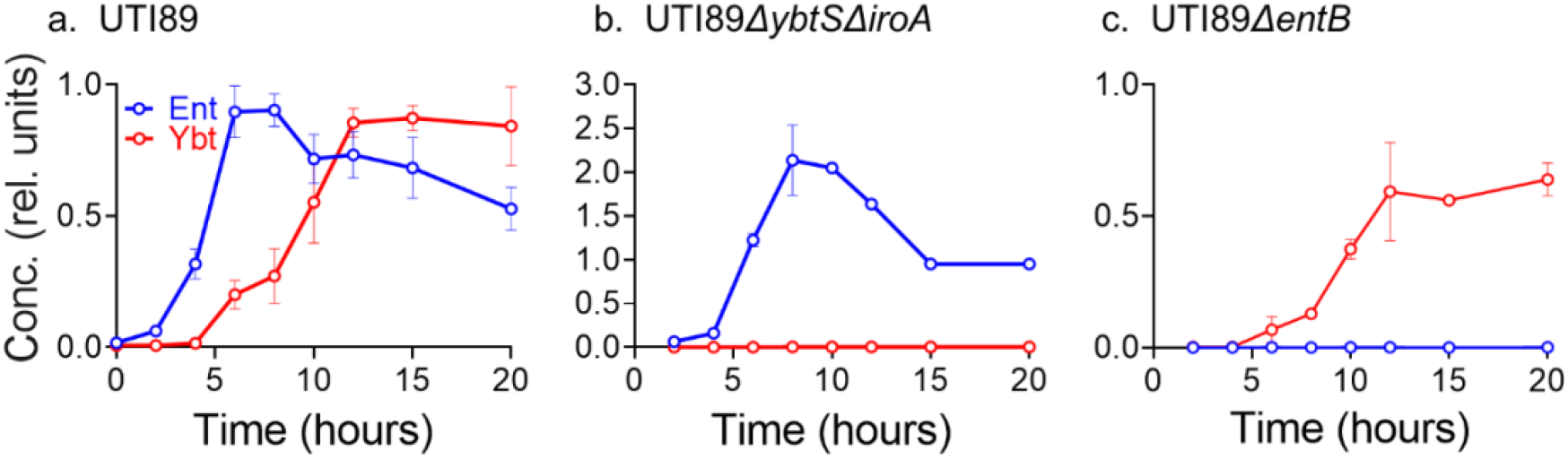
Delayed yersiniabactin production by Ent-null UTI89 is sustained in the absence of transferrin. Siderophore production by UTI89 (a) UTI89*ΔybtSΔiroA* (b), and UTI89*ΔentB* (c) in M63 medium over 20 hours of incubation. Supernatant was sampled at different times, and siderophore concentration in the conditioned medium measured by LC-MS. For each plot, the area ratio for Ent (blue) or Ybt (red) are expressed after normalization to the maximum values achieved by UTI89 for each siderophore. **a**. *p:entC-mCherry* **b**. *p:ybtP-GFP*

**Figure S4:**
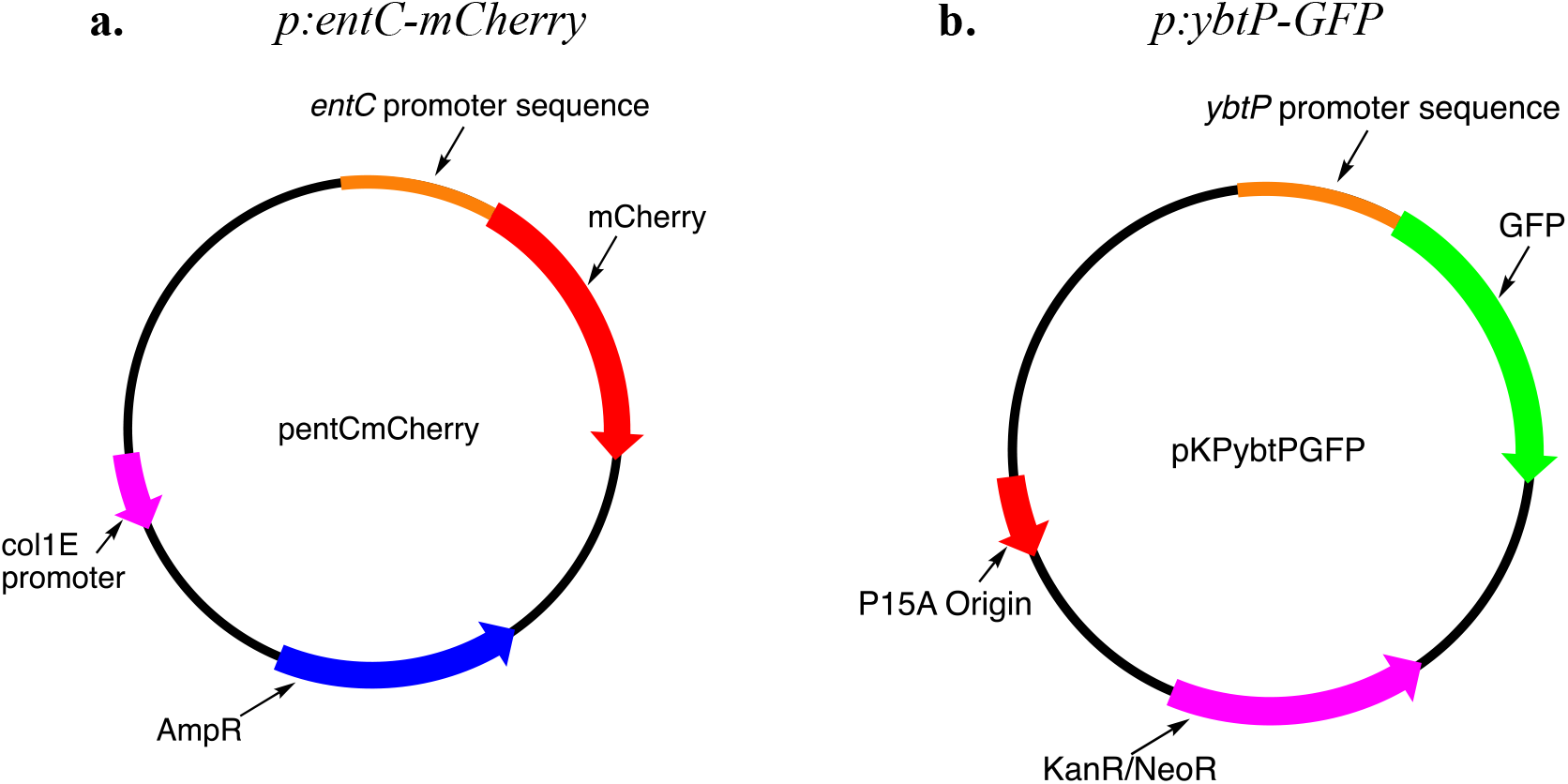
Maps of reporter plasmids containing fluorescent proteins linked to promoter sequences of operons containing early siderophore biosynthesis genes.

**Figure S5:**
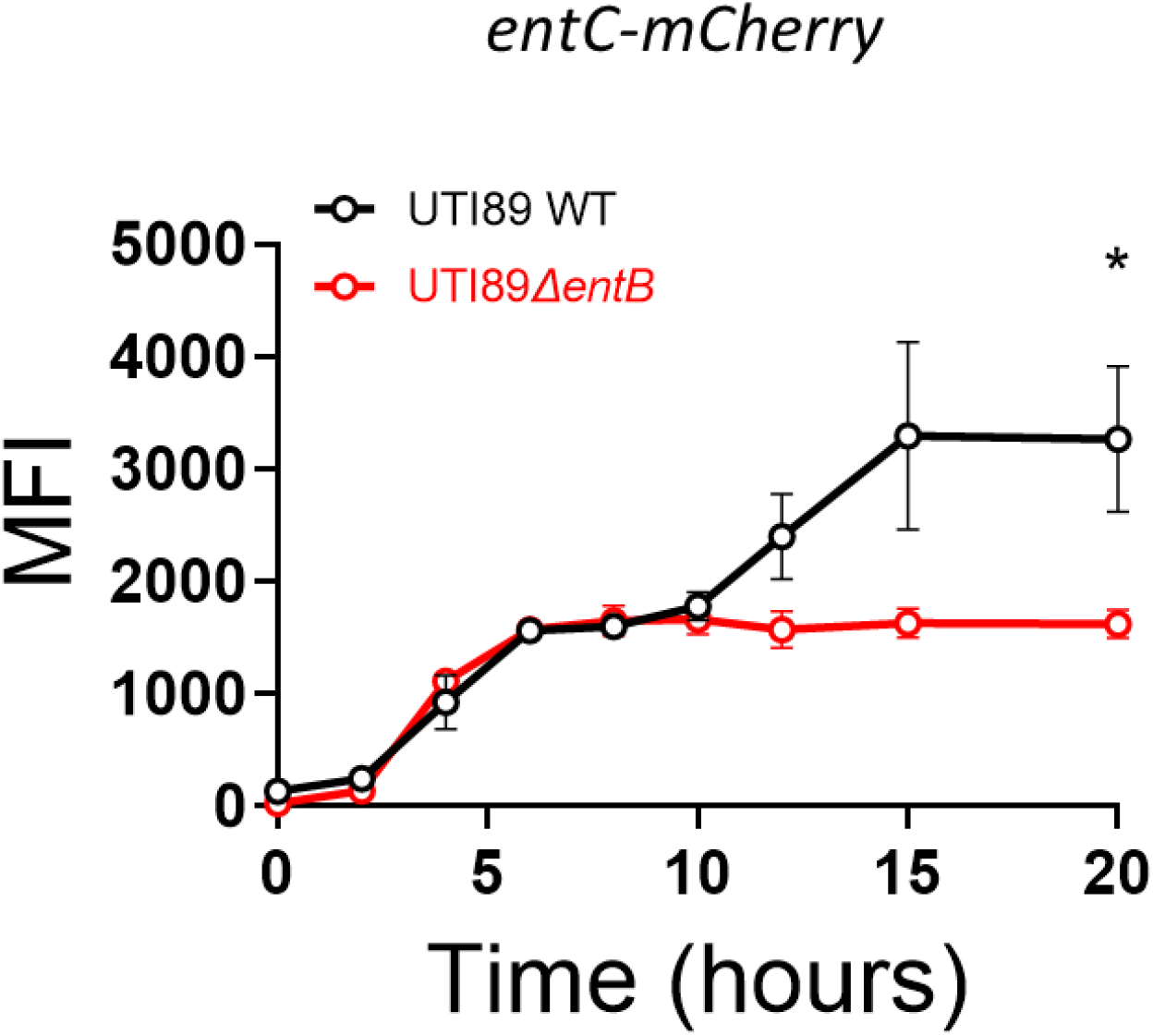
Early *entCEBA* activation is sustained in the Ent-deficient UTI89*ΔentB* mutant. UTI89 and UTI89*ΔentB* mutants transfected with the *p:entC-mCherry* and *p:ybtP-GFP* plasmids were grown in M63+hTf and expression of fluorescent proteins were measured using flow cytometry. MFI of Ent operon reporter UTI89 WT (black), UTI89*ΔentB* (red). Unpaired T-test used as the statistical test at each time point.

**Table S1:**
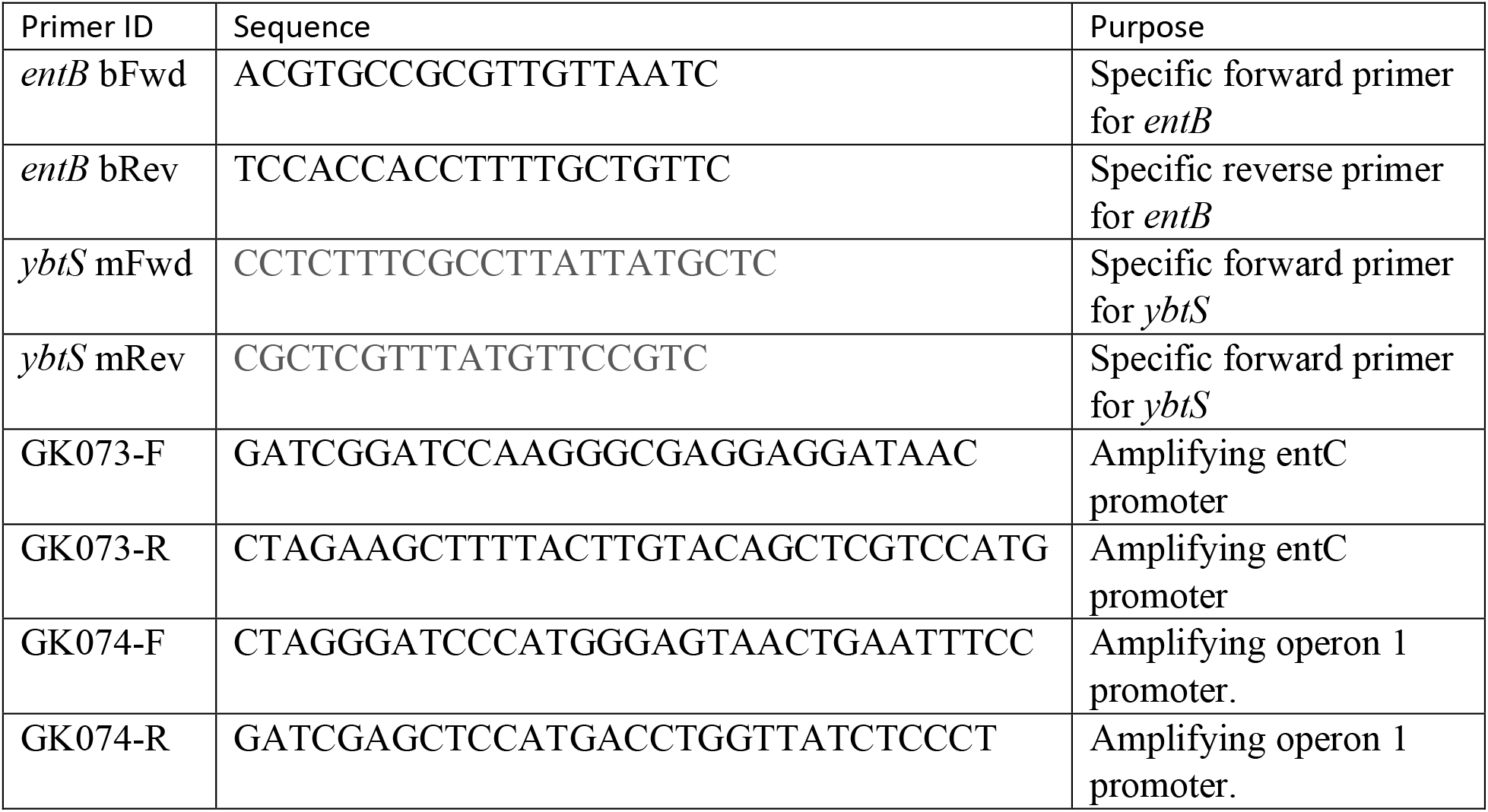
Primer sequences.

## REFERENCES

1. Foxman B. Epidemiology of urinary tract infections: incidence, morbidity, and economic costs. Am J Med. 2002;113 Suppl 1A:5S–13S. http://www.ncbi.nlm.nih.gov/pubmed/12113866. Accessed September 9, 2018.

2. Ronald A. The etiology of urinary tract infection: traditional and emerging pathogens. Dis Mon. 2003;49(2):71–82. doi:10.1067/mda.2003.8

3. Foxman B, Zhang L, Tallman P, et al. Virulence characteristics of Escherichia coli causing first urinary tract infection predict risk of second infection. J Infect Dis. 1995;172(6):1536–1541. http://www.ncbi.nlm.nih.gov/pubmed/7594713. Accessed September 9, 2018.

4. Parker KS, Wilson JD, Marschall J, Mucha PJ, Henderson JP. Network Analysis Reveals Sex- and Antibiotic Resistance-Associated Antivirulence Targets in Clinical Uropathogens. ACS Infect Dis. 2016;1(11):523–532. doi:10.1021/acsinfecdis.5b00022

5. Cassat JE, Skaar EP. Iron in Infection and Immunity. Cell Host Microbe. 2013;13(5):509. doi:10.1016/J.CHOM.2013.04.010

6. Robinson AE, Heffernan JR, Henderson JP. The iron hand of uropathogenic Escherichia coli: The role of transition metal control in virulence. Future Microbiol. 2018;13(7):813–829. doi:10.2217/fmb-2017-0295

7. Pishchany G, Haley KP, Skaar EP. Staphylococcus aureus growth using human hemoglobin as an iron source. J Vis Exp. 2013;(72):1–6. doi:10.3791/50072

8. Skaar EP, Humayun M, Bae T, DeBord KL, Schneewind O. Iron-source preference of Staphylococcus aureus infections. Science (80-). 2004;305(5690):1626–1628. doi:10.1126/SCIENCE.1099930/SUPPL_FILE/SKAAR.SOM.PDF

9. Aisen P, Leibman A, Zweier J. Stoichiometric and site characteristics of the binding of iron to human transferrin. J Biol Chem. 1978;253(6):1930–1937. doi:10.1016/s0021-9258(19)62337-9

10. Shields-cutler RR, Crowley JR, Hung CS, et al. Human urinary composition controls antibacterial activity of siderocalin. J Biol Chem. 2015;290(26):15949–15960. doi:10.1074/jbc.M115.645812

11. Shields-Cutler RR, Crowley JR, Miller CD, Stapleton AE, Cui W, Henderson JP. Human metabolome-derived cofactors are required for the antibacterial activity of siderocalin in urine. J Biol Chem. 2016;291(50):25901–25910. doi:10.1074/jbc.M116.759183

12. Correnti C, Strong RK. Mammalian siderophores, siderophore-binding lipocalins, and the labile iron pool. J Biol Chem. 2012;287(17):13524–13531. doi:10.1074/JBC.R111.311829

13. Wally J, Buchanan SK. A structural comparison of human serum transferrin and human lactoferrin. In: BioMetals. Vol 20.; 2007:249–262. doi:10.1007/s10534-006-9062-7

14. Pan Y, Sonn GA, Sin MLY, et al. Electrochemical immunosensor detection of urinary lactoferrin in clinical samples for urinary tract infection diagnosis. Biosens Bioelectron. 2010;26(2):649–654. doi:10.1016/j.bios.2010.07.002

15. Chaturvedi KS, Hung CS, Crowley JR, Stapleton AE, Henderson JP. protect pathogens during infection. Nat Chem Biol. 2012;8(8):731–736. doi:10.1038/nchembio.1020

16. Chen SL, Hung C-S, Xu J, et al. Identification of genes subject to positive selection in uropathogenic strains of Escherichia coli: a comparative genomics approach. Proc Natl Acad Sci U S A. 2006;103(15):5977–5982. doi:10.1073/pnas.0600938103

17. Hunt MD, Pettis GS, Mcintosh MA. Promoter and Operator Determinants for Fur-Mediated Iron Regulation in the Bidirectional FepA-Fes Control Region of the Escherichia Coli Enterobactin Gene System. Vol 176.; 1994. http://jb.asm.org/. Accessed March 28, 2020.

18. Fetherston JD, Bearden SW, Perry RD. YbtA, an AraC-type regulator of the Yersinia pestis pesticin/yersiniabactin receptor. Mol Microbiol. 1996;22(2):315–325. doi:10.1046/j.1365-2958.1996.00118.x

19. Christopher JP, Pistorius E, Axelrod B, et al. Ferric Uptake Regulation Protein Acts as a Repressor, Employing Iron(II) as a Cofactor To Bind the Operator of an Iron Transport Operon in Escherichia Colft. Vol 26.; 1987. https://pubs.acs.org/sharingguidelines.

20. Harris WR, Carrano CJ, Cooper SR, et al. Coordination Chemistry of Microbial Iron Transport Compounds. 19. Stability Constants and Electrochemical Behavior of Ferric Enterobactin and Model Complexes1. Am Chem Soc. 1979:6097–6104.

21. Koh EI, Henderson JP. Microbial copper-binding siderophores at the host-pathogen interface. J Biol Chem. 2015;290(31):18967–18974. doi:10.1074/jbc.R115.644328

22. Robinson AE, Lowe JE, Koh EI, Henderson JP. Uropathogenic enterobacteria use the yersiniabactin metallophore system to acquire nickel. J Biol Chem. 2018;293(39):14953–14961. doi:10.1074/jbc.RA118.004483

23. Koh EI, Robinson AE, Bandara N, Rogers BE, Henderson JP. Copper import in Escherichia coli by the yersiniabactin metallophore system. Nat Chem Biol. 2017;13(9):1016–1021. doi:10.1038/nchembio.2441

24. Henderson JP, Crowley JR, Pinkner JS, et al. Quantitative metabolomics reveals an epigenetic blueprint for iron acquisition in uropathogenic Escherichia coli. PLoS Pathog. 2009;5(2). doi:10.1371/journal.ppat.1000305

25. Zou Z, Potter RF, McCoy WH, et al. E. coli catheter-associated urinary tract infections are associated with distinctive virulence and biofilm gene determinants. JCI insight. 2023;8(2). doi:10.1172/JCI.INSIGHT.161461

26. Koh E, Hung CS, Parker KS, Crowley JR, Giblin DE, Henderson JP. Metal selectivity by the virulence-associated yersiniabactin metallophore system Eun-Ik. Metallomics. 2015;7(6):1011–1022. doi:10.1039/c4mt00341a.Metal

27. Bachman MA, Lenio S, Schmidt L, Oyler JE, Weiser JN. Interaction of Lipocalin 2, Transferrin, and Siderophores Determines the Replicative Niche of Klebsiella pneumoniae during Pneumonia. MBio. 2012;3(6):1–8. doi:10.1128/mBio.00224-11.Editor

28. Katumba GL, Tran H, Henderson JP. The Yersinia High-Pathogenicity Island Encodes a Siderophore-Dependent Copper Response System in Uropathogenic Escherichia coli. MBio. 2022;13(1). doi:10.1128/MBIO.02391-21

29. Troxell B, Hassan HM. Transcriptional regulation by Ferric Uptake Regulator (Fur) in pathogenic bacteria. Front Cell Infect Microbiol. 2013;3(OCT). doi:10.3389/FCIMB.2013.00059

30. Teixidó L, Carrasco B, Alonso JC, Barbé J, Campoy S. Fur Activates the Expression of Salmonella enterica Pathogenicity Island 1 by Directly Interacting with the hilD Operator In Vivo and In Vitro. PLoS One. 2011;6(5):19711. doi:10.1371/JOURNAL.PONE.0019711

31. Delany I, Rappuoli R, Scarlato V. Fur functions as an activator and as a repressor of putative virulence genes in Neisseria meningitidis. Mol Microbiol. 2004;52(4):1081–1090. doi:10.1111/J.1365-2958.2004.04030.X

32. García-Lara J, Shang LH, Rothfield LI. An extracellular factor regulates expression of sdiA, a transcriptional activator of cell division genes in Escherichia coli. J Bacteriol. 1996;178(10):2742–2748. doi:10.1128/JB.178.10.2742-2748.1996

33. Sperandio V, Torres AG, Girón JA, Kaper JB. Quorum sensing is a global regulatory mechanism in enterohemorrhagic Escherichia coli O157:H7. J Bacteriol. 2001;183(17):5187–5197. doi:10.1128/JB.183.17.5187-5197.2001

34. Yang S, Lopez CR, Zechiedrich EL. Quorum sensing and multidrug transporters in Escherichia coli. Proc Natl Acad Sci U S A. 2006;103(7):2386–2391. doi:10.1073/PNAS.0502890102/SUPPL_FILE/02890FIG6.PDF

35. Wang L, Li J, March JC, Valdes JJ, Bentley WE. luxS-dependent gene regulation in Escherichia coli K-12 revealed by genomic expression profiling. J Bacteriol. 2005;187(24):8350–8360. doi:10.1128/JB.187.24.8350-8360.2005/ASSET/03550AA2-7A4D-49CA-BCF1-A426E540423F/ASSETS/GRAPHIC/ZJB0240553010007.JPEG

36. McRose DL, Baars O, Seyedsayamdost MR, Morel FMM. Quorum sensing and iron regulate a two-for-one siderophore gene cluster in Vibrio harveyi. Proc Natl Acad Sci U S A. 2018;115(29):7581–7586. doi:10.1073/pnas.1805791115

37. Lewenza S, Sokol PA. Regulation of ornibactin biosynthesis and N-acyl-L-homoserine lactone production by cepR in Burkholderia cepacia. J Bacteriol. 2001;183(7):2212–2218. doi:10.1128/JB.183.7.2212-2218.2001/ASSET/12A0813A-D644-4575-8FEE-D848E1708B51/ASSETS/GRAPHIC/JB0711270003.JPEG

38. Lewenza S, Conway B, Greenberg EP, Sokol PA. Quorum sensing in Burkholderia cepacia: identification of the LuxRI homologs CepRI. J Bacteriol. 1999;181(3):748–756. doi:10.1128/JB.181.3.748-756.1999

39. Wen Y, Kim IH, Son JS, Lee BH, Kim KS. Iron and Quorum Sensing Coordinately Regulate the Expression of Vulnibactin Biosynthesis in Vibrio vulnificus. J Biol Chem. 2012;287(32):26727. doi:10.1074/JBC.M112.374165

40. Anisimov R, Brem D, Heesemann J, Rakin A. Transcriptional regulation of high pathogenicity island iron uptake genes by YbtA. Int J Med Microbiol. 2005;295(1):19–28. doi:10.1016/J.IJMM.2004.11.007

41. Fetherston JD, Bearden SW, Perry RD. YbtA, an AraC-type regulator of the Yersinia pestis pesticin/yersiniabactin receptor. Mol Microbiol. 1996;22(2):315–325. doi:10.1046/J.1365-2958.1996.00118.X

42. Hultgren SJ, Schwan WR, Schaeffer AJ, Duncan JL. Regulation of production of type 1 pili among urinary tract isolates of Escherichia coli. Infect Immun. 1986;54(3):613–620. doi:10.1128/IAI.54.3.613-620.1986

43. Ohlemacher SI, Giblin DE, D’Avignon DA, Stapleton AE, Trautner BW, Henderson JP. Enterobacteria secrete an inhibitor of Pseudomonas virulence during clinical bacteriuria. J Clin Invest. 2017;127(11):4018–4030. doi:10.1172/JCI92464

44. Eickhoff MJ, Fei C, Cong JP, Bassler BL. LuxT Is a Global Regulator of Low-Cell-Density Behaviors, Including Type III Secretion, Siderophore Production, and Aerolysin Production, in Vibrio harveyi. MBio. 2022;13(1). doi:10.1128/MBIO.03621-21/SUPPL_FILE/MBIO.03621-21-ST001.DOCX

45. Stintzi A, Evans K, Meyer JM, Poole K. Quorum-sensing and siderophore biosynthesis in Pseudomonas aeruginosa: lasR/lasI mutants exhibit reduced pyoverdine biosynthesis. FEMS Microbiol Lett. 1998;166(2):341–345. doi:10.1111/J.1574-6968.1998.TB13910.X

46. Lewenza S, Sokol PA. Regulation of ornibactin biosynthesis and N-acyl-L-homoserine lactone production by CepR in Burkholderia cepacia. J Bacteriol. 2001;183(7):2212–2218. doi:10.1128/JB.183.7.2212-2218.2001

47. Scholz RL, Greenberg EP. Sociality in Escherichia coli: Enterochelin is a private good at low cell density and can be shared at high cell density. J Bacteriol. 2015;197(13):2122–2128. doi:10.1128/JB.02596-14

48. Mysorekar IU, Hultgren SJ. Mechanisms of uropathogenic Escherichia coli persistence and eradication from the urinary tract. Proc Natl Acad Sci U S A. 2006;103(38):14170–14175. doi:10.1073/PNAS.0602136103/SUPPL_FILE/02136FIG10.PDF

49. Chaturvedi KS, Hung CS, Giblin DE, et al. Cupric yersiniabactin is a virulence-associated superoxide dismutase mimic. ACS Chem Biol. 2014;9(2):551–561. doi:10.1021/cb400658k

50. Schubert S, Rakin A, Heesemann J. The Yersinia high-pathogenicity island (HPI): evolutionary and functional aspects. Int J Med Microbiol. 2004;294(2-3):83–94. doi:10.1016/J.IJMM.2004.06.026

51. Lukacik P, Barnard TJ, Keller PW, et al. Structural engineering of a phage lysin that targets Gram-negative pathogens. Proc Natl Acad Sci U S A. 2012;109(25):9857–9862. doi:10.1073/PNAS.1203472109/-/DCSUPPLEMENTAL

52. Gao H, Zhou D, Li Y, et al. The iron-responsive fur regulon in Yersinia pestis. J Bacteriol. 2008;190(8):3063–3075. doi:10.1128/JB.01910-07

53. Perry RD, Fetherston JD. Yersinia pestis--etiologic agent of plague. Clin Microbiol Rev. 1997;10(1):35. doi:10.1128/CMR.10.1.35

54. Guo C, Steinberg LK, Cheng M, Song JH, Henderson JP, Gross ML. Site-Specific Siderocalin Binding to Ferric and Ferric-Free Enterobactin As Revealed by Mass Spectrometry. ACS Chem Biol. 2020;15(5):1154–1160. doi:10.1021/ACSCHEMBIO.9B00741

55. Schwyn B, Neilands JB. Universal chemical assay for the detection and determination of siderophores. Anal Biochem. 1987;160(1):47–56. doi:10.1016/0003-2697(87)90612-9

56. Chen SL, Hung C-S, Xu J, et al. Identification of genes subject to positive selection in uropathogenic strains of Escherichia coli: a comparative genomics approach. Proc Natl Acad Sci U S A. 2006;103(15):5977–5982. doi:10.1073/pnas.0600938103

57. Lv H, Henderson JP. Yersinia high pathogenicity Island genes modify the Escherichia coli primary metabolome independently of siderophore production. J Proteome Res. 2011;10(12):5547–5554. doi:10.1021/pr200756n

